# Delay-Induced Uncertainty in Physiological Systems

**DOI:** 10.1101/2020.07.17.209544

**Authors:** Bhargav Karamched, George Hripcsak, Dave Albers, William Ott

## Abstract

Medical practice in the intensive care unit is based on the supposition that physiological systems such as the human glucose-insulin system are *reliabile*. Reliability of dynamical systems refers to response to perturbation: A dynamical system is reliable if it behaves predictably following a perturbation. Here, we demonstrate that reliability fails for an archetypal physiological model, the Ultradian glucose-insulin model. Reliability failure arises because of the presence of delay. Using the theory of rank one maps from smooth dynamical systems, we precisely explain the nature of the resulting delay-induced uncertainty (DIU). We develop a recipe one may use to diagnose DIU in a general dynamical system. Guided by this recipe, we analyze DIU emergence first in a classical linear shear flow model and then in the Ultradian model. Our results potentially apply to a broad class of physiological systems that involve delay.

## 1. Introduction

Delay can significantly impact the dynamics of physiological systems at multiple scales. At the level of genetic regulatory networks, distributed delay on the order of minutes results from the transcriptional, translational, and post-translational steps that result in the production of functional regulator proteins. Such delay can accelerate signaling within feedforward architectures [27], alter the statistics of noise-induced switching phenomena [19, 33], and produce oscillations in synthetic genetic circuits [57]. This paper is about a novel route through which delay can cause reliability loss within concrete dynamical systems of interest in physiology and biomedicine. We call the resulting unreliability *delay-induced uncertainty* (DIU). We believe that DIU has profound implications for oscillations that arise in systems physiology, especially the ultradian glucose-insulin oscillation observed within human endocrine physiology.

The concept of reliability plays a central role in control theory [29, 45, 53, 56, 60, 61, 79]. Here, we adopt the following conceptual definition: A dynamical system is *reliable* if system response to perturbation does not depend (after a transient) on the state of the system at the time of perturbation. In this paper, we specifically consider external forcing signals that perturb a system’s intrinsic dynamics. This notion of reliability has been used to study phase oscillator networks subjected to Brownian inputs [37]. Here, we use concepts from ergodic theory to make our conceptual definition mathematically precise. In particular, we leverage geometric and analytical ideas from the theory of nonuniformly hyperbolic dynamical systems and specifically the theory of rank one maps [67, 69, 70].

Clinical and laboratory practice throughout biology, physiology, and medicine proceeds from the assumption that the dynamics of measured quantities are reliable. For instance, a clinician administers medication to a patient based on the belief that the medical intervention will not induce an unpredictable, erratic response. While chaotic dynamics have been observed in some physiological models [1, 16, 17, 34, 41], reliability failure in physiology is not yet well-understood by mathematicians, nor is its significance known to clinicians. It is vital to uncover the mechanisms that produce reliability loss and develop precise mathematical characterizations of the resulting dynamics. It is vital to assess the impact of reliability loss on data assimilation and clinical practice. This paper uncovers one such mechanism.

We formulate a general recipe for the emergence of DIU in dynamical systems and then focus on damped, driven oscillators. The recipe consists of three ingredients. First, delay renders the unforced dynamical system excitable. For the damped, driven oscillators we consider in this paper, delay produces a weakly stable limit cycle. This can happen, for instance, via a delay-induced supercritical Hopf bifurcation [7, 28, 54, 71, 75, 76]. Second, the unforced system possesses intrinsic shear. For damped, driven oscillators, shear quantifies velocity gradients near the limit cycle. Third, the forcing drive interacts with the shear to create hyperbolicity in the dynamics. This hyperbolicity causes the damped, driven oscillator to lose reliability. Geometrically, the interaction between the shear and the forcing drive stretches and folds the phase space, producing positive Lyapunov exponents.

Why is DIU important in physiological settings? We approach this question by analyzing an important physiological model, the glucose-insulin Ultradian model [15, 30, 59]. The Ultradian system was originally constructed as a model with the minimal number of components needed to explain ultradian oscillations. Delayed regulatory feedback between glucose and insulin produces the oscillation. Although there are other models with more expansive explanations of ultradian oscillations, the Ultradian model has been used to accurately describe and predict glucose dynamics in humans [3]. Nevertheless, the *reliability* of ultradian oscillations has not yet been studied. In this paper, we carefully demonstrate that DIU emerges in the Ultradian system.

The presence of DIU in glucose-insulin dynamics may have profound implications for clinical care in the intensive care unit (ICU), where glucose and insulin treatments (external forcing drives) are central to glycemic management. More generally, DIU is potentially relevant for any physiological system wherein delayed regulatory feedback controls try to maintain healthy homeostasis. From the clinical perspective, the presence of DIU in the Ultradian model implies that even if we know the overall health state (model parameters) of the patient exactly, the result of a therapeutic intervention can nevertheless depend sensitively on the location of the patient trajectory in phase space at the time of the intervention. This sensitivity may well confuse the clinician, since the same intervention might *succeed at some times and fail at others*.

We organize the scientific content of this paper as follows. We begin by contextualizing DIU within mathematical physiology and biomedicine. In particular, we explain why we have selected the Ultradian model for DIU demonstration. We then describe our DIU recipe and use it to show that DIU emerges in a delay variant of the linear shear flow model first studied by Zaslavsky [77] and then by Lin and Young [38]. The delay linear shear flow setting explains the mathematics and geometry that underpins DIU. Our results for delay linear shear flow establish that the infinite-dimensional dynamical systems (flows on function spaces) generated by delay differential equations can produce DIU. We then turn to the Ultradian model, for which we perform a variety of experiments that establish and probe DIU. Our results for both models are numerical in nature and use the maximal Lyapunov exponent as a reliability diagnostic. When applicable, we use the theory of rank one maps [67,69,70] to rigorously support our numerical findings. We conclude the paper by discussing open mathematical questions inspired by DIU, as well as the importance of DIU in biomedicine and clinical practice.

## 2. Contextualizing DIU: Mathematical physiology and clinical care

DIU is about the interplay between intrinsic delay dynamics (mathematical physiology) and external forcing drives (clinical care). Both can be complex dynamical processes [2, 11, 25, 26, 52].

### 2.1. Ultradian model

In this paper we have elected to demonstrate the presence of DIU in the Ultradian model, a minimal glucose-insulin model [2,59] that includes glucose-insulin dynamics, several feedback mechanisms that represent insulin-mediated glucose regulation by the pancreas [59], and hepatic responses. The Ultradian model has successfully described human glucose dynamics in controlled laboratory settings. It has been well-validated, and is used in practice [3, 59].

The Ultradian model is a compartment model devised to explain observed ultradian oscillations. It was built by researchers who added model subsystems and states until ultradian oscillatory dynamics emerged. They found that two compartments describing insulin, namely plasma and interstitial insulin, should be present, along with the appropriate coupling. Further, they reasoned that a delayed feedback between insulin secretion and glucose released by the liver should be included to produce ultradian oscillations. Other hypotheses regarding sources of ultradian oscillations exist [39], but they all rely on delays of some kind and still include the delayed hepatic response.

We focus on the Ultradian model because it is the simplest model with ultradian oscillations. We believe our conclusions will hold for more complex models, such as models with multiple sources of delay.

### 2.2. Brief survey of the glucose-insulin modeling landscape

Since the creation of the minimal model [6], glucose-insulin modeling has branched in many directions. There exist models for the ICU setting [22, 31, 32, 35], such as the ICU minimal model (ICUMM) [23, 65], the ICING model [36], and models that can potentially resolve ICU-related issues such as stress hypoglycemia [20, 21, 39, 51, 59, 63] or the source of ultradian glycemic oscillations [39, 59].

Models have been created to capture type-2 diabetes pathogenesis in particular [20,21,63]. Such models may apply directly to the oral and intraveneous glucose tolerance test settings (IVGTT, OGTT) [10, 20], or to both GTT and free-living settings [20, 43]. The interplay between type-2 diabetes and digestion has been assessed [12]. Stochastic differential equation models have been designed to robustly handle the complexities of clinical data [43].

Endocrine models have been designed to apply to patients with type-1 diabetes as well [5, 42, 44, 49, 50, 72, 74], but this family of models is qualitatively distinct from the others. In particular, type-1 diabetes implies no endogenous insulin production, and therefore no glycemic oscillations are induced by the automatic control system of the body.

### 2.3. Processes of glycemic management

In this paper, we establish the presence of DIU for glucose signaling in the Ultradian model. That said, we expect DIU to be relevant for general forcing drives, interpreted here as mathematical representations of clinical care. Clinical care can be complex and dynamic. We highlight this here by describing two scenarios in detail, namely glycemic management in the ICU and type-2 diabetes self-management. These scenarios are consistent with the fact that the Ultradian model includes endogenous insulin. For each scenario, we describe the external forcing (clinical intervention methods) here, and then speculate about DIU impact in Section 6.

In the ICU, clinicians begin by maximizing the amount of nutrition, delivered enterally (via a tube), because critical care patients are almost never able to consume enough nutrition. Delivered nutrition is perturbed with some frequency because some interventions require cessation of nutrition; furthermore, the amount of nutrition a patient can tolerate often drifts because of changes in health state or other interventions [2]. Blood glucose levels may elevate for a variety of reasons, including stress, intervention response, and other unknown mechanisms. Exogenously delivered insulin manages elevated glucose levels. Summarizing, both exogenous insulin and nutrition vary over time.

Moreover, the health care system itself also impacts glycemic management in the ICU, and should therefore be accounted for when modeling glucose management. In particular, the current standard of care for glycemic management at our institutions calls for nurses to determine insulin administration using a complex flowsheet based on the current glucose target range and insulin infusion. A diversity of such non-personalized protocols are in use [73].

Type-2 diabetes, a chronic and often life-long disease, is managed by the patient in an outpatient setting [3,13,14]. Management usually includes a mix of diet, exercise, and medications. Exogenous insulin is only used for relatively severe cases. Note that while nutrition is delivered continuously in the ICU, type-2 patients must deal with pulsatile nutritional drivers (meal consumption). Management usually amounts to measuring glucose with a finger stick, often in the morning, and before and after meals. These measurements inform decisions such as what and how much to eat and when and how to exercise. Summarizing, type-2 diabetes management involves temporally-localized kicks [2] (nutrition consumption) along with exercise perturbations and insulin administration.

### 2.4. Scope of DIU impact

While we focus on glucose-insulin dynamics here, the use of mathematical physiology within medicine has broad potential [4, 78]. DIU is relevant for any system that features delay and oscillatory dynamics. Such systems show up throughout physiology. Examples include pulmonary and respiratory dynamics [41, 55], cardiac dynamics [8], female endocrine dynamics [18, 64], and neurological dynamics [9, 24, 58], to name but a few. It will be important to assess the existence and impact of DIU in these contexts, as DIU may confound our current understanding of these systems.

## 3. Mathematics of delay-induced uncertainty

We consider delay dynamical systems subjected to external forcing drives. On an intuitive level, we say that such a system is reliable if system response to external stimuli (after a transient period) does not depend on the state of the system at the time of stimulus initiation. We say that delay-induced uncertainty (DIU) occurs when reliability fails. This paper focuses on a specific route to DIU.

### 3.1. DIU recipe

Our route to DIU involves the following ingredients. We describe them intuitively here and later follow with precise mathematical formulations.

#### (U1) Delay-induced excitability

Delay renders the unforced (intrinsic) dynamical system excitable. This happens, for instance, when delay produces a weakly stable invariant structure (stationary point, homoclinic orbit, limit cycle). As we will see, delay in the Ultradian model produces a limit cycle via a supercritical Hopf bifurcation.

#### (U2) Intrinsic shear

When the unforced system possesses a limit cycle, shear refers to significant velocity gradients near the limit cycle. Atmospheric wind shear provides a good mental picture of the phenomenon.

#### (U3) External forcing allows shear to act

External forcing allows the shear to stretch and fold the phase space, thereby creating hyperbolicity in the dynamics.

Importantly, DIU is not a phenomenon wherein the external forcing simply overwhelms the intrinsic dynamics. Forcing amplitudes can be quite small. In the DIU framework, forcing acts as an amplifier, amplifying intrinsic shear to produce rich, complex dynamics.

We now examine a concrete linear model, delay linear shear flow, in detail. This examination illuminates the geometry and dynamics of DIU.

### 3.2. Delay linear shear flow

This archetypal system allows us to describe what happens when systems with limit cycles experience pulsatile forcing drives. The dynamics take place on the cylinder 𝕊^1^ × ℝ. Writing *θ* for the 𝕊^1^-coordinate and *z* for the ℝ-coordinate, delay linear shear flow is generated by the delay differential equations

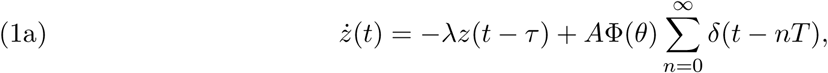

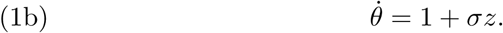

System (1) is infinite-dimensional due to the delay *τ* : One must specify an initial history *h* : [− *τ*, 0] → 𝕊^1^ ×ℝ in order to propagate the dynamics forward. The final term on the right side of Eq. (1a) represents periodic pulsatile kicks. Here *A* ⩾ 0 is the kick amplitude, *T* > 0 is the inter-kick time, Φ : 𝕊^1^ → ℝ describes the kick profile, and *δ* is the Dirac delta. The Dirac delta has the following interpretation: At each nonnegative integer multiple of *T*, each point in 𝕊^1^ × ℝ instantaneously moves from (*θ, z*) to (*θ, z* + *A*Φ(*θ*)). Between kicks, the dynamics are governed by the unforced delay equations

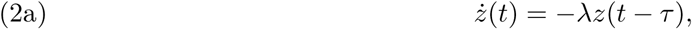

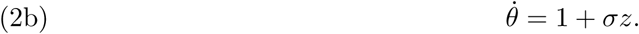

Together with delay *τ*, the parameters *λ* > 0 and *σ* ∈ ℝ shape the dynamics of the unforced dynamical system.

#### The delay-free case *τ* = 0

In the absence of delay, linear shear flow assumes the form

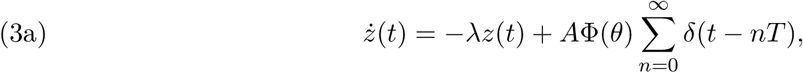

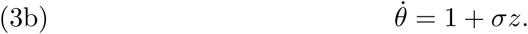

This deceptively simple system has been studied by mathematical physicists such as Zaslavsky [77] and by dynamicists such as Lin and Young [38]. Without forcing (*A* = 0), the set Ω = {(*θ, z*) : *z* = 0} is a limit cycle of system (3) for all values of the contraction parameter *λ* > 0. This limit cycle is weakly stable if *λ* is small. The dynamics of linear shear flow depend on the size and shape of the pulsatile kicks, as well as how trajectories relax to the limit cycle between kicks.

The angular velocity gradient parameter *σ* quantifies the shear in system (3). This parameter links with *A* and *λ* to form the *hyperbolicity factor*

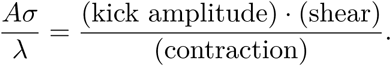

The hyperbolicity factor governs the dynamics of system (3). We express this through the behavior of the time-*T* map *F*_*T*_ : 𝕊^1^ ×ℝ → 𝕊^1^ ×ℝ generated by linear shear flow (the result of one kick followed by one relaxation cycle).

If 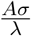 is small, then the kicked system will quickly return to equilibrium after each kick (Figure 1a). Mathematically, *F*_*T*_ admits an attractor diffeomorphic to 𝕊^1^. However, stretch and fold geometry emerges when the hyperbolicity factor is large (Figure 1b). Provided the kick profile Φ is not a constant function, each kick creates wave-like variability in the *z*-direction (Figure 1b, first image). If the relaxation time *T* is sizable, shear will then cause the waves to stretch and fold (Figure 1b, second image). Chaotic behavior consequently emerges when the hyperbolicity factor is large, chaos that is both sustained in time and observable.

**Figure 1.**
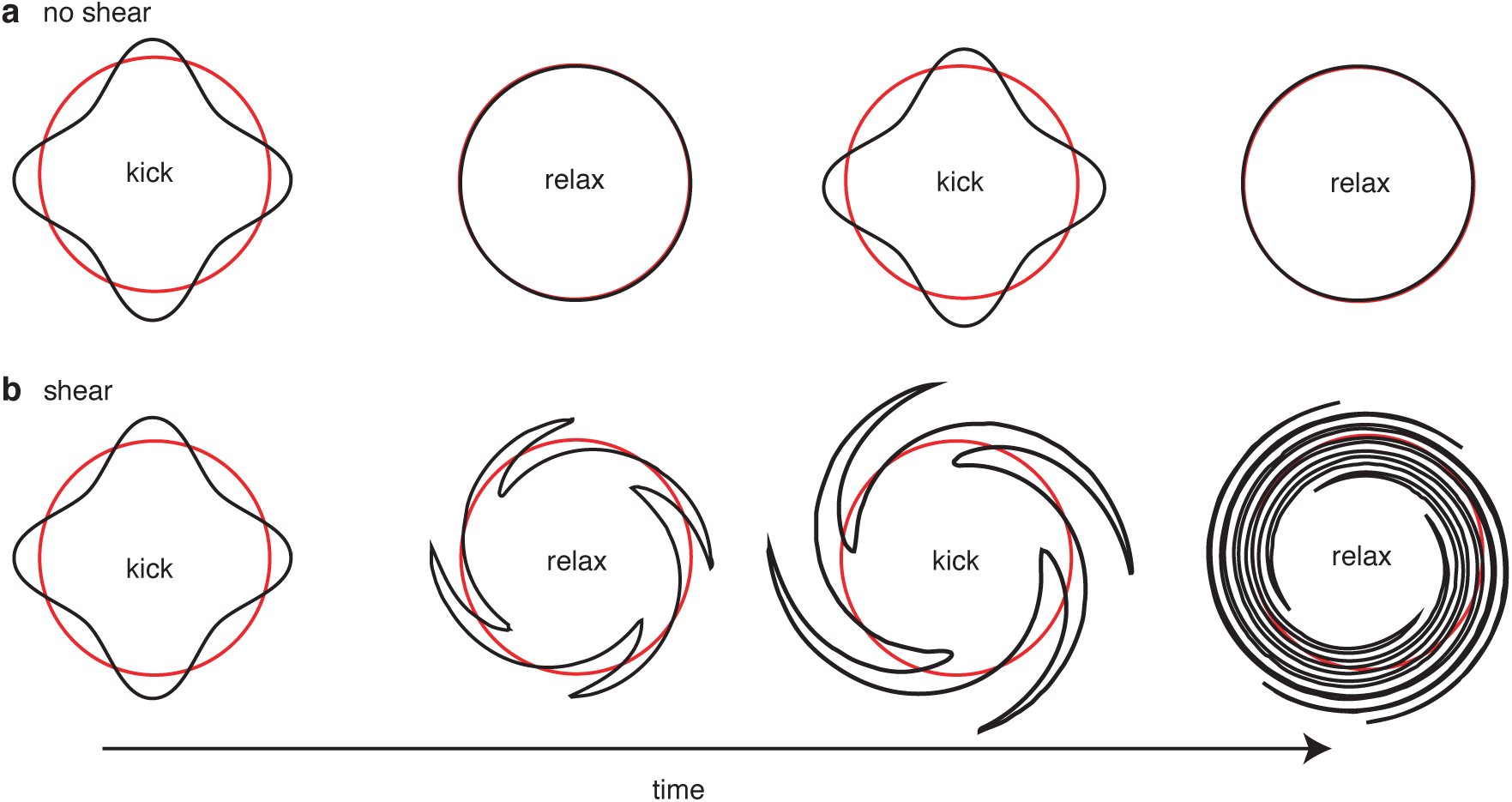
Kick-relaxation dynamics of linear shear flow (3). When the system is kicked, the limit cycle (red circle) deforms (black curve). Before the next kick, the system relaxes toward the limit cycle. **(a)** When the hyperbolicity factor 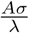 is small, the kicked limit cycle quickly relaxes. In this case, the phase space does not stretch and fold. The time-*T* map *F*_*T*_ admits an attractor that is diffeomorphic to 𝕊^1^. **(b)** When the hyperbolicity factor is large, the kicked limit cycle stretches and folds during relaxation (assuming the relaxation time *T* is large enough to allow stretch and fold geometry to manifest). The kick-relaxation cycle produces sustained, observable chaos characterized by the theory of rank one maps.

Mathematically, *F*_*T*_ admits a strange attractor for a set of *T* values of positive Lebesgue measure, provided the hyperbolicity factor is sufficiently large and the kick profile Φ is not a constant function. For such values of *T, F*_*T*_ is genuinely nonuniformly hyperbolic. In particular, it exhibits a positive Lyapunov exponent (the chaos is sustained in time). The strange attractor supports a unique ergodic Sinai-Ruelle-Bowen (SRB) measure. This measure describes the asymptotic distribution of almost every orbit in the basin of attraction (the chaos is observable). The SRB measure possesses strong statistical properties: The system obeys a dynamical version of the central limit theorem, correlations for Hölder observable decay exponentially, and a large deviations principle holds.

Found in [68], precise statements and proofs of these rigorous results for linear shear flow rely on the theory of rank one maps. Developed by Wang and Young [67,69,70], rank one theory provides a platform for proving the existence of nonuniformly hyperbolic dynamics (sustained, observable chaos) within concrete systems of interest in the biochemical and physical sciences.

The proofs for linear shear flow involve verifying the hypotheses of the theory of rank one maps. In particular, Wang and Young [68] analyze a certain ‘infinite relaxation’ *T* → ∞ limit of *F*_*T*_. This procedure produces the singular limit, a parametrized family of circle maps. Rank one theory links the dynamics of the singular limit to those of *F*_*T*_. For linear shear flow, the hyperbolicity factor gives the amount of expansion in the singular limit. Expansion in the singular limit links to sustained, observable chaos for *F*_*T*_.

#### The delay case *τ* > 0

We argue that DIU emerges when *τ* > 0. Since no theory of rank one maps exists yet for delay systems, we compute the top Lyapunov exponent Λ_max_ numerically and use Λ_max_ as a DIU diagnostic. That is, DIU is present if Λ_max_ > 0, whereas the system is reliable if Λ_max_ *<* 0. To argue that DIU emerges, we establish the (U1)–(U3) route to DIU.

(U1). Delay induces excitability in system (1) because the stability of the limit cycle weakens as *τ* increases. This is because the strength of stability of the zero solution to the scalar delay differential equation

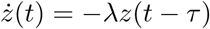

is determined by the (complex) solutions of the characteristic equation

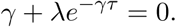

(U2). Shear is present in system (1) exactly as it is present in delay-free linear shear flow because the 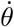 equation does not depend on *τ*.

(U3). Here there exists a subtle difference between the delay case and the delay-free case. In the delay-free case, the kick profile Φ needs to be nonconstant in order to create the *z*-variability that leads to stretching and folding of the phase space. This requirement can be dropped in the delay case. Here, the history *h* that serves as initial data for system (1) can provide *z*-variability, so DIU can emerge even if Φ is a constant function. Indeed, we demonstrate this in Figure 2.

**Figure 2.**
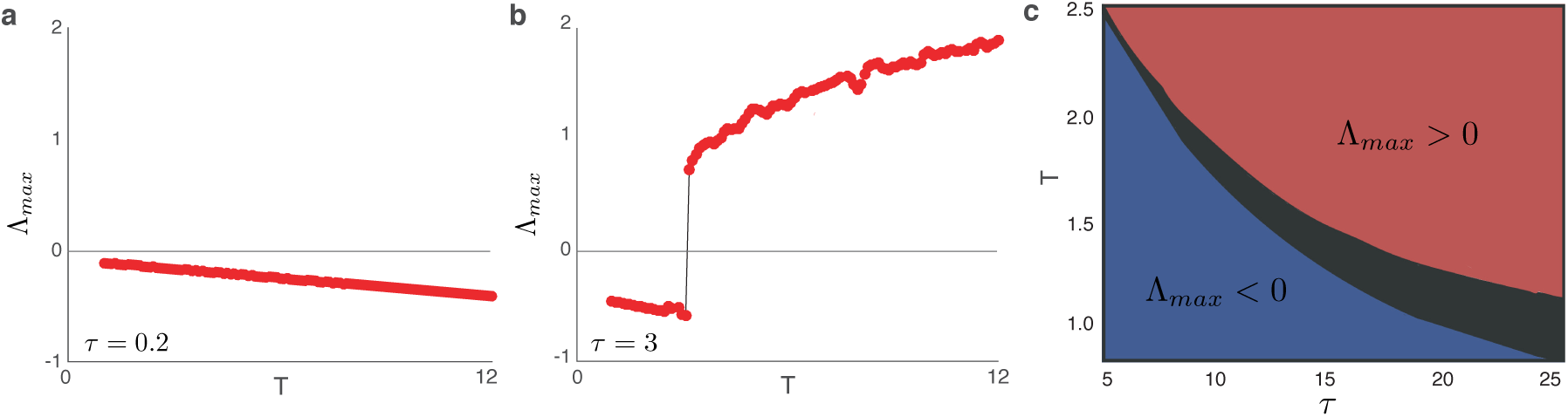
DIU for delay linear shear flow (1). **(a)** When delay *τ* is small, the top Lyapunov exponent Λ_max_ is negative over the tested range of inter-kick times *T*. DIU has not emerged when delay is small. **(b)** When delay is substantial, DIU emerges when Λ_max_ transitions from negative to positive. **(c)** Heatmap illustrating the sign of Λ_max_ as a function of the delay *τ* and the inter-kick time *T*. Red indicates that Λ_max_ > 0; blue indicates that Λ_max_ *<* 0. The sign of Λ_max_ depends sensitively on *τ* and *T* in the black region. Here, *σ* = 3, *λ* = 0.1, *A* = 0.1, and Φ = 1. For *t* ∈ [−*τ*, 0], we take *h*(*t*) = (*θ*(*t*), *z*(*t*)) = (0, *t*^2^).

### 3.3. Nonlinear systems

Rigorous results analogous to those we have desribed for delay-free linear shear flow exist for finite-dimensional nonlinear systems as well. Importantly, the weakly stable invariant structure referred to in (U1) can be a stationary point [46], a homoclinic orbit [66], or a limit cycle [47]. In each case, shear must be identified and carefully quantified. All of these invariant structures are relevant from the physiological point of view.

We focus in the present work on the weakly stable limit cycle that emerges via supercritical Hopf bifurcation. This setting has been analyzed for finite-dimensional nonlinear systems [68] and for certain parabolic evolution partial differential equations [40]. We now present convincing evidence that for the Ultradian model of glucose-insulin dynamics, DIU emerges in the wake of a delay-induced Hopf bifurcation.

## 4. The Ultradian model

We now investigate DIU in a canonical glucose-insulin model commonly referred to as the Ultradian model [15, 30, 59]. The Ultradian model is a compartment model with three state variables: plasma glucose (*G*), plasma insulin (*I*_*p*_), and interstitial insulin (*I*_*i*_), see Fig. 3. These three state variables are coupled to a three-stage linear delay filter, producing a 6-dimensional phase space. The Ultradian model is particularly popular because it is the simplest physiological model that captures the main features of glucose-insulin oscillations [15,59] and provides a mechanistic description of the cause of the oscillations. The model includes two major negative feedback loops describing effects of insulin on glucose use and glucose production, and both loops include glucose-based stimulation of insulin secretion. Oscillations in the model depend on (i) a time delay of 30-45 minutes for the effect of insulin on glucose production and (ii) the slow effect of insulin on glucose use arising from insulin being in two distinct compartments. We focus on the former in this paper. Note that the Ultradian model includes explicit physiological delay, but the system is *finite-dimensional* because the delay assumes the form of a three-stage linear filter.

**Figure 3.**
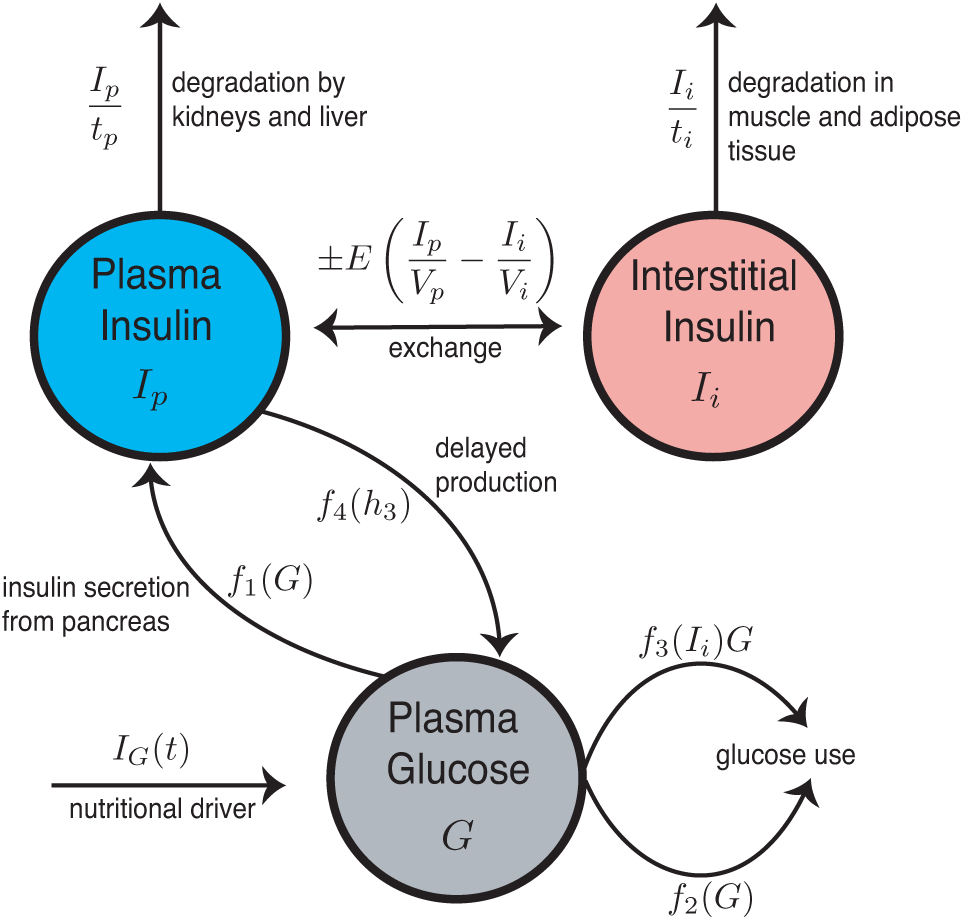
Schematic of the Ultradian model of glucose-insulin dynamics.

The full model is given by

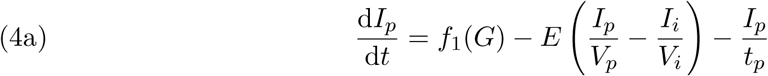

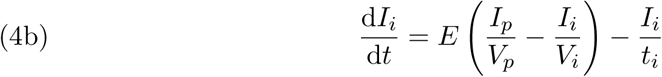

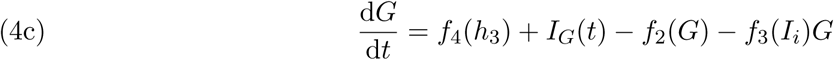

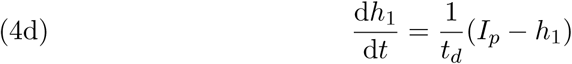

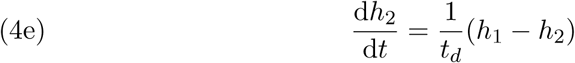

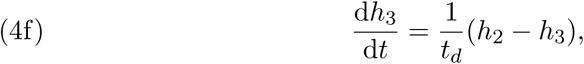

where *f*_1_(*G*) represents the rate of insulin production, *f*_2_(*G*) represents insulin-independent glucose use, *f*_3_(*I*_*i*_)*G* represents insulin-dependent glucose use, and *f*_4_(*h*_3_) represents delayed insulin-dependent glucose use. The functional forms of these are

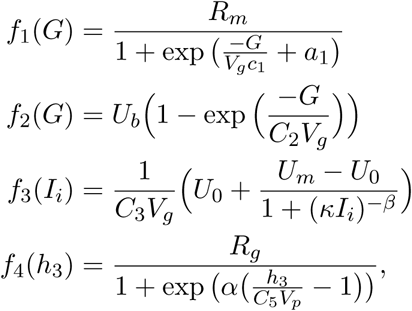

with

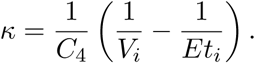

See Table 1 for parameter descriptions and nominal parameter values.

**Table 1.** Full list of parameters for the Ultradian glucose-insulin model [3]. Note that IIGU and IDGU denote insulin-independent glucose utilization and insulin-dependent glucose utilization, respectively.

| Ultradian model parameters |  |  |
| --- | --- | --- |
| Name | Nominal Value | Meaning |
| $V_p$ | 3 L | plasma volume |
| $V_i$ | 11 L | interstitial volume |
| $V_g$ | 10 L | glucose space |
| $E$ | $0.2 \text{ L min}^{-1}$ | exchange rate for insulin between remote and plasma compartments |
| $t_p$ | 6 min | time constant for plasma insulin degradation (via kidney and liver filtering) |
| $t_i$ | 100 min | time constant for remote insulin degradation (via muscle and adipose tissue) |
| $t_d$ | 12 min | delay between plasma insulin and glucose production |
| $R_m$ | $209 \text{ mU min}^{-1}$ | linear constant affecting insulin secretion |
| $a_1$ | 6.6 | exponential constant affecting insulin secretion |
| $C_1$ | $300 \text{ mg L}^{-1}$ | exponential constant affecting insulin secretion |
| $C_2$ | $144 \text{ mg L}^{-1}$ | exponential constant affecting IIGU |
| $C_3$ | $100 \text{ mg L}^{-1}$ | linear constant affecting IDGU |
| $C_4$ | $80 \text{ mU L}^{-1}$ | factor affecting IDGU |
| $C_5$ | $26 \text{ mU L}^{-1}$ | exponential constant affecting IDGU |
| $U_b$ | $72 \text{ mg min}^{-1}$ | linear constant affecting IIGU |
| $U_0$ | $4 \text{ mg min}^{-1}$ | linear constant affecting IDGU |
| $U_m$ | $94 \text{ mg min}^{-1}$ | linear constant affecting IDGU |
| $R_g$ | $180 \text{ mg min}^{-1}$ | linear constant affecting IDGU |
| $\alpha$ | 7.5 | exponential constant affecting IDGU |
| $\beta$ | 1.772 | exponent affecting IDGU |

We consider an idealized nutritional driver *I*_*G*_(*t*) that includes a basal signal and pulsatile kicks. The nutritional driver is given by

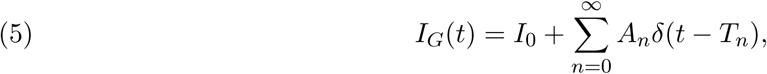

where *I*_0_ is a basal nutritional input for the system, *T*_*n*_ is the time of the *n*th feeding, and *A*_*n*_ is the amount of carbohydrate in that meal. The signal *I*_*G*_(*t*) represents the external forcing in the Ultradian model. The form of *I*_*G*_(*t*) in Eq. (5) produces the following dynamics: Between two consecutive kicks (*T*_*n* −1_ *< t < T*_*n*_), Ultradian dynamics evolve according to system (4) with *I*_*G*_(*t*) = *I*_0_. At the time *T*_*n*_ of meal *n*, the glucose state variable, *G*, undergoes the instantaneous change *G* ↦ *G* + *A*_*n*_.

We investigate the emergence of DIU for several different models of kick amplitudes 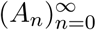 and inter-kick times 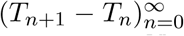. In reality, meals produce glucose kicks that are temporally localized but not instantaneous. Nevertheless, our DIU results for the idealized nutritional driver strongly predict DIU emergence for the complex external forcing drives encountered in the intensive care unit.

## 5. DIU in the Ultradian model

We deploy our DIU recipe to establish DIU emergence in the Ultradian model. We show that DIU emerges for both constant and random kick amplitudes and inter-kick times. The results of this section are numerical in nature for two reasons. First, system (4) is difficult to analyze because of the many nonlinearities. Even computing equilibria for the unforced system (obtained by removing the nutritional driver *I*_*G*_(*t*)) requires solving transcendental equations. More importantly, though, the theory of rank one maps has not yet been extended to treat random inter-kick times or random kick amplitudes. Even when both kick amplitude and inter-kick time are held constant (*A*_*n*_ = *A* and *T*_*n*_ = *nT* for all *n*), the current theory of rank one maps cannot tell us if a strange attractor exists for *specific* values of *A* and *T* [47, 68]. Rather, the rigorous applications of the theory developed thus far prove the existence of strange attractors for parameter sets of positive Lebesgue measure.

We therefore analyze the Ultradian model numerically and use the maximal Lyapunov exponent, Λ_max_, as a DIU diagnostic: Λ_max_ > 0 indicates DIU, while Λ_max_ *<* 0 indicates reliability. We make connections with the theory of rank one maps whenever possible.

### 5.1. Parameter selection

Excluding the nutritional driver, we set all Ultradian model parameters to the values in Table 1 for our simulations. For the nutritional driver, we set the basal rate *I*_0_ to zero. We are therefore free only to tune the delay *t*_*d*_ and choose models for the kick amplitudes *A*_*n*_ and the inter-kick times *T*_*n*+1_ − *T*_*n*_.

### 5.2. The DIU recipe for the Ultradian model

As we did with delay linear shear flow, we verify (U1)–(U3) for the Ultradian system.

(U1). Consider the unforced version of system (4), obtained by removing *I*_*G*_(*t*). For a variety of time delays, Figure 4 shows glucose timeseries and two-dimensional projections of phase space trajectories generated by the unforced Ultradian system. We see that a stable equilibrium bifurcates into a weakly stable limit cycle as the system undergoes a supercritical Hopf bifurcation at a delay value 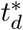 satisfying 8 < 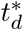 < 12. The presence of the weakly stable Hopf limit cycle implies excitability once the supercritical Hopf bifurcation has occurred.

**Figure 4.**
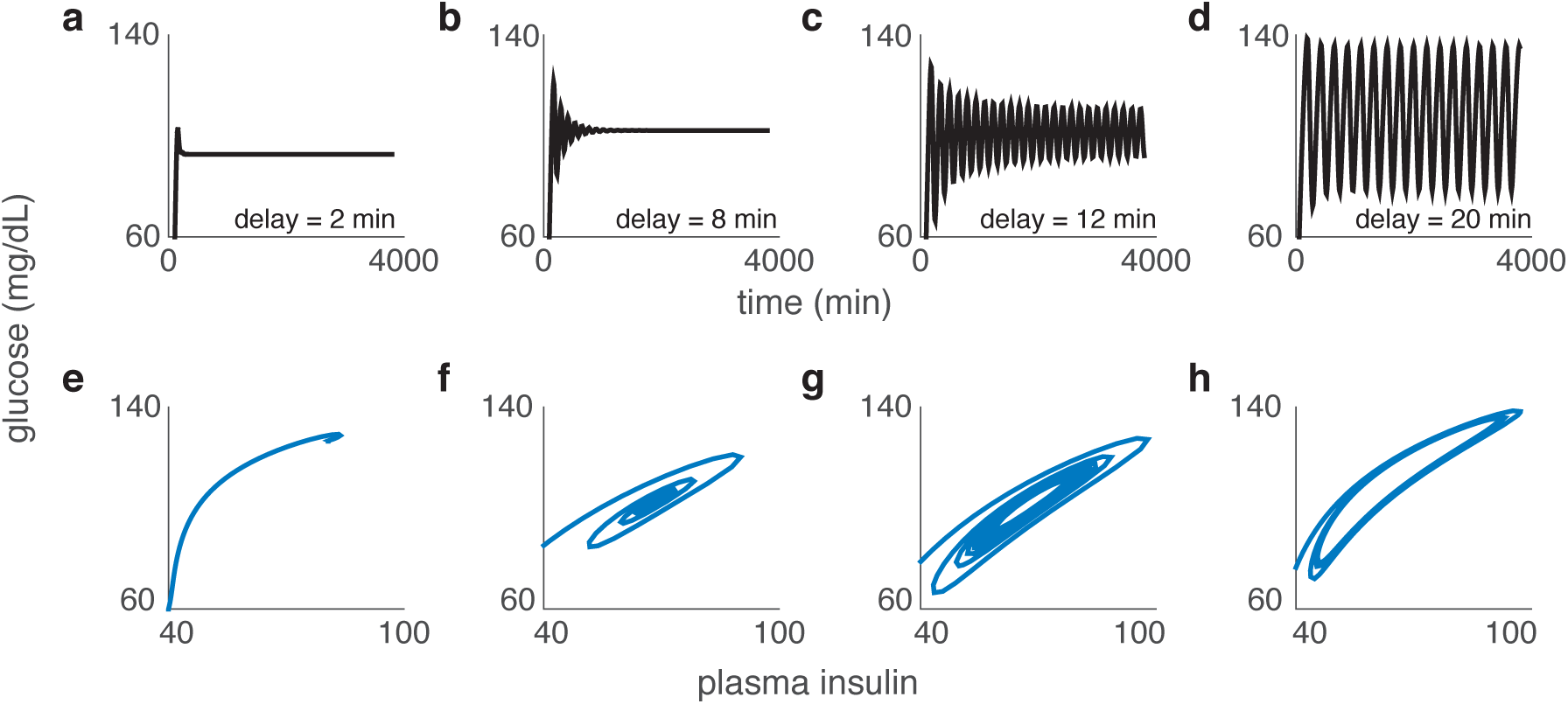
As delay *t*_*d*_ increases, a supercritical Hopf bifurcation produces a limit cycle in the unforced Ultradian model. Top row: Glucose timeseries for **(a)** *t*_*d*_ = 2 min, **(b)** *t*_*d*_ = 8 min, **(c)** *t*_*d*_ = 12 min, and **(d)** *t*_*d*_ = 20 min. Bottom row: Projection of a phase space trajectory onto glucose-plasma insulin space for **(e)** *t*_*d*_ = 2 min, **(f)** *t*_*d*_ = 8 min, **(g)** *t*_*d*_ = 12 min, and **(h)** *t*_*d*_ = 20 min. Other parameter values are given in Table 1.

(U2) and (U3). We claim that limit cycles subjected to pulsatile forcing drives generically satisfy these two recipe ingredients. The geometric ideas of Wang and Young [68] and the quantitative analysis of Ott and Stenlund [47] support this claim for finite-dimensional nonlinear systems. For (U2), shear can be understood geometrically by examining the shape of the strong stable foliation in a tubular neighborhood of the limit cycle [68]. Ott and Stenlund [47] quantify shear by defining a *shear integral* that represents the accumulation of shear as one traverses the limit cycle.

For (U3), the external forcing must interact with the shear in order to produce strange attractors. We claim that this happens for generic pulsatile forcing drives. To support this claim, Wang and Young prove that given a *C*^4^ flow on a Riemannian manifold that admits a hyperbolic limit cycle, periodic kicks will produce strange attractors for an open set of *C*^3^ kick functions (Theorem 1 of [68]). Ott and Stenlund [47] define a function that quantifies the forcing-shear interaction and assume that this function is Morse in their main theorem on the existence of strange attractors. They conjecture that this assumption will hold for a generic kick-generating vector field, both in terms of topological genericity and prevalence. See Remark 2.1 of [47] for a discussion of the conjecture and [48] for information about prevalence. Note that for the Ultradian model, the kicks provided by the nutritional driver have no spatial variation *with respect to the original state variables*. Such variation should be present, however, *for the coordinate system near the limit cycle* developed in [47].

We now present the numerical experiments that establish the presence of DIU in the Ultradian model. We use the maximal Lyapunov exponent as a DIU diagnostic: Λ_max_ > 0 indicates DIU, while Λ_max_ *<* 0 indicates reliability.

We compute the maximal Lyapunov exponent by solving system (4) as follows. During the relaxation intervals (*T*_*n* −1_, *T*_*n*_) between kicks, we integrate the differential equations using MATLAB’s ode23s stiff solver. At kick times *T*_*n*_, we pause the differential equation solver and apply the diffeomorphism of phase space induced by the kick *G* ↦ *G* + *A*_*n*_. We compute Λ_max_ by completing 10^5^ relaxation-kick cycles.

#### Constant kick amplitude, periodic or Poissonian kicks

For our first set of experiments, we choose a value of the delay *t*_*d*_ such that the Hopf limit cycle is present in the unforced system, and then hold *t*_*d*_ fixed. We consider kicks of constant intensity, *A*_*n*_ = *A* for all *n*. Kick times are either periodic, *T*_*n*_ = *nT* for all *n*, or Poissonian. In the Poissonian case, the inter-kick times *T*_*n*+1_ − *T*_*n*_ are independent and exponentially distributed with mean *T*. We show that DIU emerges even for these relatively simple forms of the nutritional driver (5) by examining how the maximal Lyapunov exponent depends on *A* and *T*.

We compute the maximal Lyapunov exponent as follows. For simulations involving periodic kicks (see Figs. 5a-c, 6a-c), we track two solutions to system (4), separated at time *t* = 0 by an amount *d*_0_ = 10^−8^. Think of one of these solutions as the base solution and the other as a secondary, perturbative solution. After completing the first relaxation window (0, *T*), we apply the glucose kick *G* ↦ *G* + *A* at time *T*. After the kick, we compute the separation between the solutions at time *T*, denoted *d*_1_, and store the quantity log(*d*_1_*/d*_0_) in a vector. We then renormalize by moving the secondary orbit toward the base orbit so that the distance between the two resets to *d*_0_. We proceed in this manner for 10^5^ relaxation-kick cycles of the form (*nT*, (*n* + 1)*T*]. This produces a vector containing 10^5^ values of log(*d*_1_*/d*_0_). Averaging over this vector produces Λ_max_, the maximal Lyapunov exponent.

**Figure 5.**
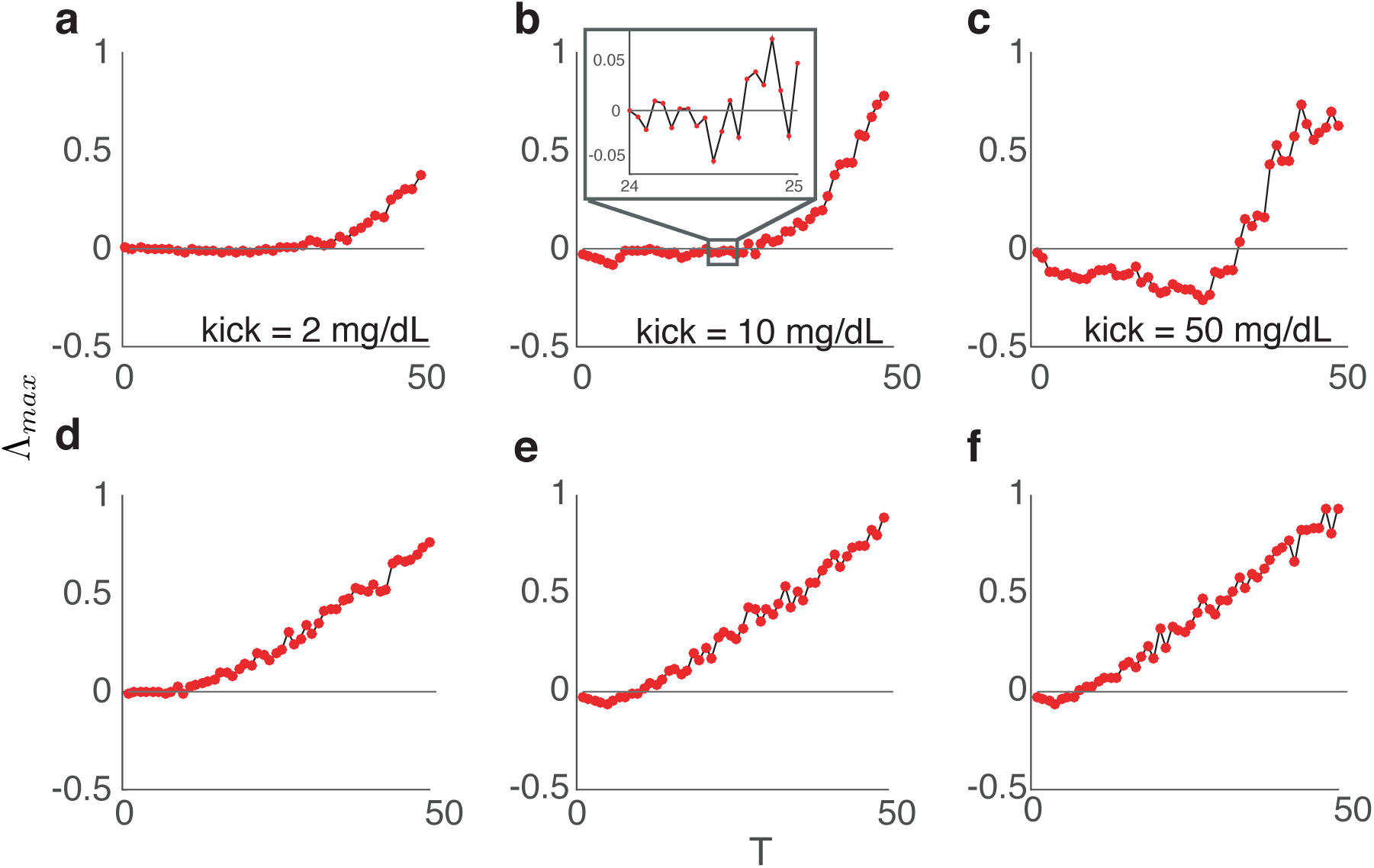
Maximal Lyapunov exponent in the Ultradian model as a function of kick timing. Positive Λ_max_ indicates the presense of DIU. **(a)-(c)** For periodic kicks, plots of Λ_max_ versus the inter-kick time *T* for several fixed values of kick amplitude *A*. **(a)** *A* = 2 mg/dL. **(b)** *A* = 10 mg/dL. **(c)** *A* = 50 mg/dL. **(d)-(f)** For Poissonian kicks, plots of Λ_max_ versus mean inter-kick time *T* for several fixed values of kick amplitude *A*. **(d)** *A* = 2 mg/dL. **(e)** *A* = 10 mg/dL. **(f)** *A* = 50 mg/dL. Other parameter values are as in Table 1.

For simulations involving Poissonian kicks (see Figs. 5d-f, 6d-f), the maximal Lyapunov exponent is a random variable, as it depends *a priori* on the random inter-kick times. To compute it, we first sample 10^5^ inter-kick times from the exponential distribution with mean *T*. These samples produce a single realization of the stochastic process. We compute the maximal Lyapunov exponent for this realization by proceeding as we did in the case of periodic kicks. That is, we compute log(*d*_1_*/d*_0_) following each relaxation-kick cycle and then average. Finally, we average the realization-dependent maximal Lyapunov exponent over 10^5^ realizations of the Poisson process. Abusing notation slightly, we call this average Λ_max_.

Figures 5a-c and 6a-c display maximal Lyapunov exponent results for the case of constant kick amplitude and periodic kicks. Here, Λ_max_ is a function of the kick amplitude *A* and the inter-kick time *T*.

For three different fixed values of *A*, Λ_max_ becomes positive as *T* increases, indicating the onset of DIU (Figure 5a-c). This is consistent with the intuition gained from analyzing the linear shear flow model: Larger values of *T* allow more time for the phase space to stretch and fold between kicks. The maximal Lyapunov exponent depends on *T* in a particularly interesting way when *A* = 50 mg/dL (Figure 5c). Here, Λ_max_ *<* 0 for small values of *T*, indicating that the system is reliable and suggesting that the time-*T* map of the system possesses an attractor that is diffeomorphic to the limit cycle present in the unforced Ultradian system. By constrast, Λ_max_ > 0 for large values of *T*, indicating the presence of DIU and suggesting that the time-*T* map of the system possesses a strange attractor. The inset in Figure 5b shows that fluctuates around zero for moderately large values of *T*. This suggests that the time-*T* map of the system possesses horseshoes (transient chaos).

In Figure 6a-c, we compute Λ_max_ as a function of *A* for different fixed values of *T*. When *T* is small (*T* = 5 min), the maximal Lyapunov exponent is negative for all of the values of *A* we have simulated, indicating that the system is reliable, robustly so with respect to *A*, when *T* is small (Figure 6a). By contrast, when *T* is large (*T* = 100 min), Λ_max_ is positive for all of the values of *A* we have tested, indicating that DIU is present even when *A* is small (Figure 6c). We observe a transition from reliability to DIU as *A* increases when *T* is moderately large (*T* = 20 min, Figure 6b).

**Figure 6.**
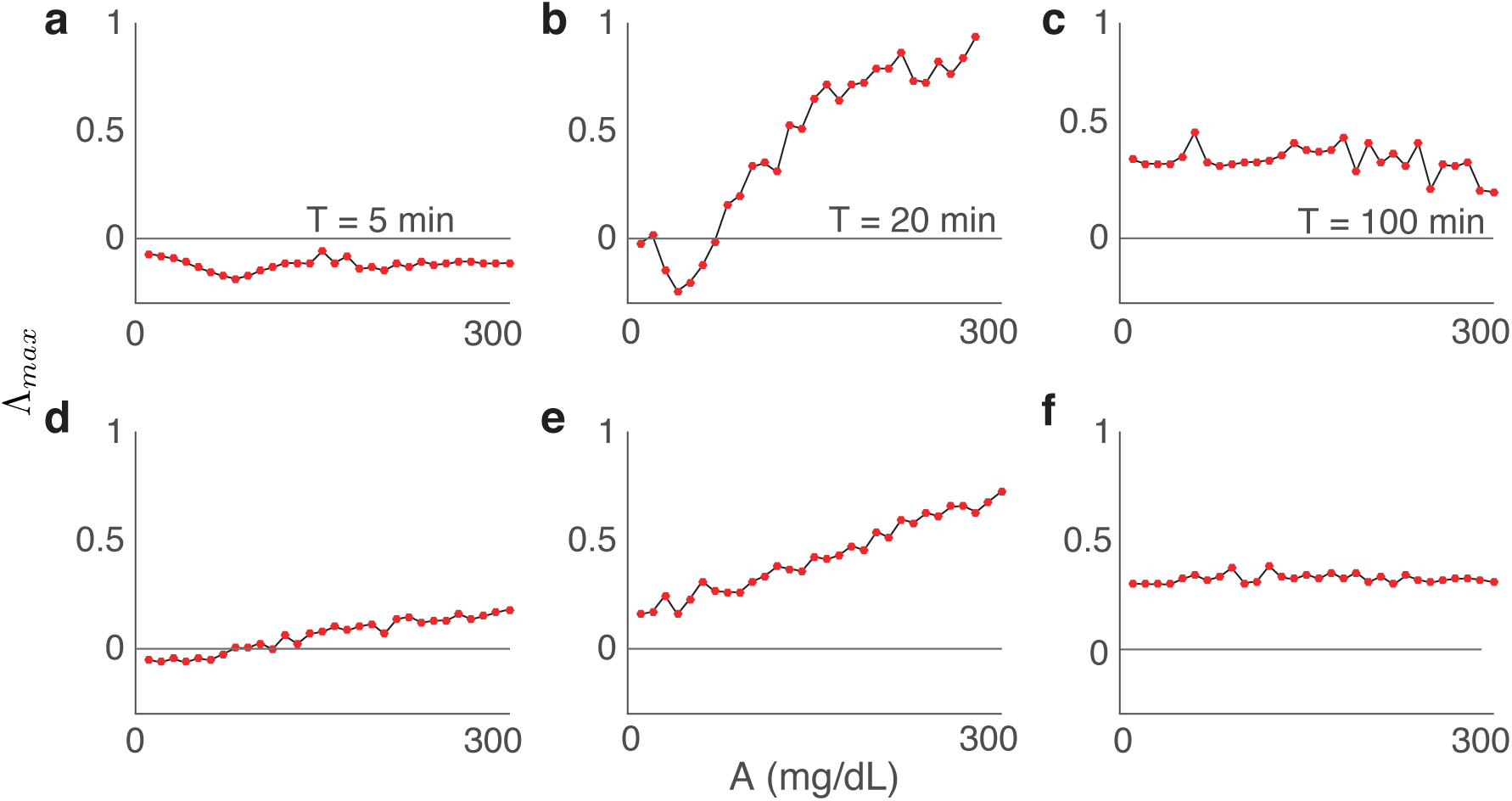
Maximal Lyapunov exponent in the Ultradian model as a function of kick amplitude. Positive Λ_max_ indicates the presense of DIU. **(a)-(c)** For periodic kicks, plots of Λ_max_ as a function of kick amplitude for several different values of inter-kick time *T*. **(a)** *T* = 5 min. **(b)** *T* = 20 min. **(c)** *T* = 100 min. For Poissonian kicks, plots of Λ_max_ as a function of kick amplitude for several different values of mean inter-kick time *T*. **(d)** *T* = 5 min. **(e)** *T* = 20 min. **(f)** *T* = 100 min. Other parameter values are as in Table 1.

Figures 5d-f and 6d-f display maximal Lyapunov exponent results for the case of constant kick amplitude and Poissionian kicks. Here, Λ_max_ > 0 for most of the kick amplitudes *A* and mean inter-kick times *T* we tested, indicating robust presence of DIU. Note that when *A* is fixed at *A* = 50 mg/dL, DIU onset occurs significantly earlier in the Poissonian case than in the periodic case as *T* increases (Figure 5c,f).

Figure 7 illustrates how the Ultradian dynamics differ in the reliable case and the DIU case. Here, we simulate the Ultradian model with delay *t*_*d*_ = 12 min, a value for which the unforced system possesses a limit cycle. We drive the system with periodic glucose kicks of constant amplitude *A* = 10 mg/dL. We select a small value of the inter-kick time for which the forced system behaves reliably (*T* = 20 min), and a larger value for which the system exhibits DIU (*T* = 200 min). The first two rows of Figure 7 show representative glucose timeseries and corresponding empirical glucose distributions for the reliable case (left column) and the DIU case (right column). In the reliable case, glucose levels oscillate regularly as expected, but interestingly the empirical glucose distribution is bimodal. By contrast, the erratic behavior of the glucose timeseries in the DIU case reflects the chaos in the system. Notice that the empirical distribution in the DIU case appears to be approximately Gaussian. This observation is consistent with the results on SRB measures for general kicked limit cycles in [47, 68], where scalar observables are shown to satisfy a dynamical version of the central limit theorem.

**Figure 7.**
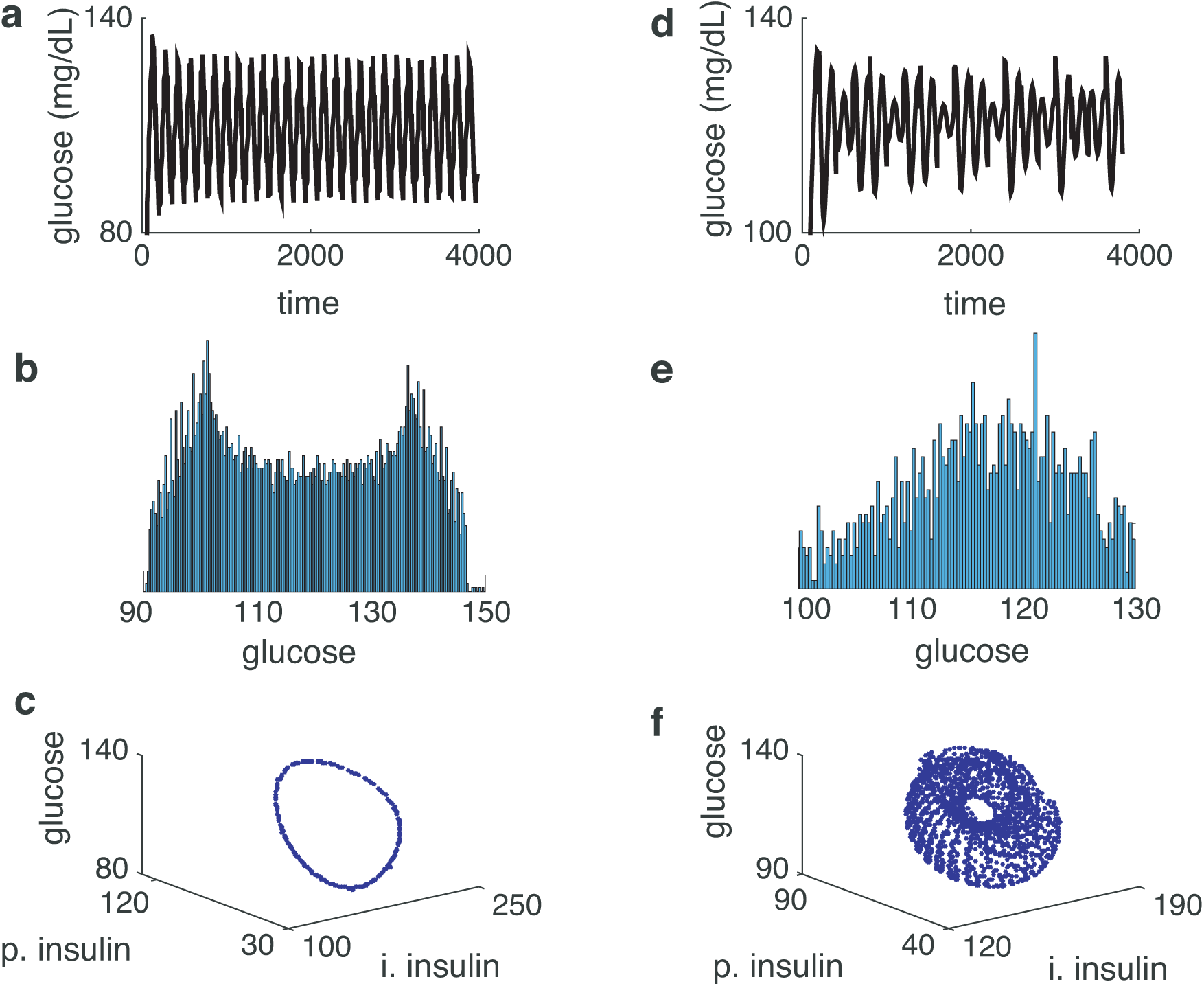
Dynamical profile of Ultradian dynamics in the reliable case (left column) and the DIU case (right column). **(a)** Glucose levels oscillate in a regular manner. **(b)** The empirical glucose distribution associated with the timeseries from (a) is bimodal. **(c)** The time-*T* map generated by the Ultradian system possesses an attractor that is diffeomorphic to the limit cycle of the unforced system. **(d)** Glucose levels evolve erratically. **(e)** The empirical glucose distribution associated with the timeseries from (d) appears to be approximately Gaussian. **(f)** The time-*T* map generated by the Ultradian system possesses a strange attractor. Left column: *T* = 20 min. Right column: *T* = 200 min. All panels: *t*_*d*_ = 12 min, *A* = 10 mg/dL, and other parameter values are as in Table 1.

Figure 7d,e suggests that the impact of DIU on clinical practice will be subtle and complex. We have framed the present paper in terms of reliability. Framed this way, we associate DIU with reliability loss, a clear potential problem when attempting rational medical intervention. But reliability takes a *single-orbit perspective* on dynamics. If only the *statistical* behavior of observables of the dynamics (such as glucose level) matters in a particular setting, then DIU may be *beneficial*, since the results of [47, 68] suggest that observables of Ultradian dynamics behave with a high level of statistical regularity.

We plot the attractors of the time-*T* map generated by the Ultradian system in Figure 7c,f. In the reliable case, the attractor is diffeomorphic to the limit cycle of the unforced system. In the DIU case, we observe a strange attractor with intricate geometry. These results are consistent with the rigorous theory of [47, 68].

#### Uniformly distributed kick amplitudes, periodic kicks

For this set of experiments, we make the (more realistic) assumption that kick amplitudes are random, rather than constant. In particular, we assume the kick amplitudes *A*_*n*_ are independent and uniformly distributed, while the kicks are periodic in time with inter-kick time *T*. Figure 8a shows the distribution of Λ_max_ as a function of *T* when the kick amplitudes are drawn from the uniform distribution on [45, 55]. When *T* is small, the distribution of Λ_max_ is essentially a Dirac delta at a negative value. Interestingly, at the moment *E*[Λ_max_] crosses zero, the variance of Λ_max_ immediately becomes positive, and continues to grow as *T* increases.

**Figure 8.**
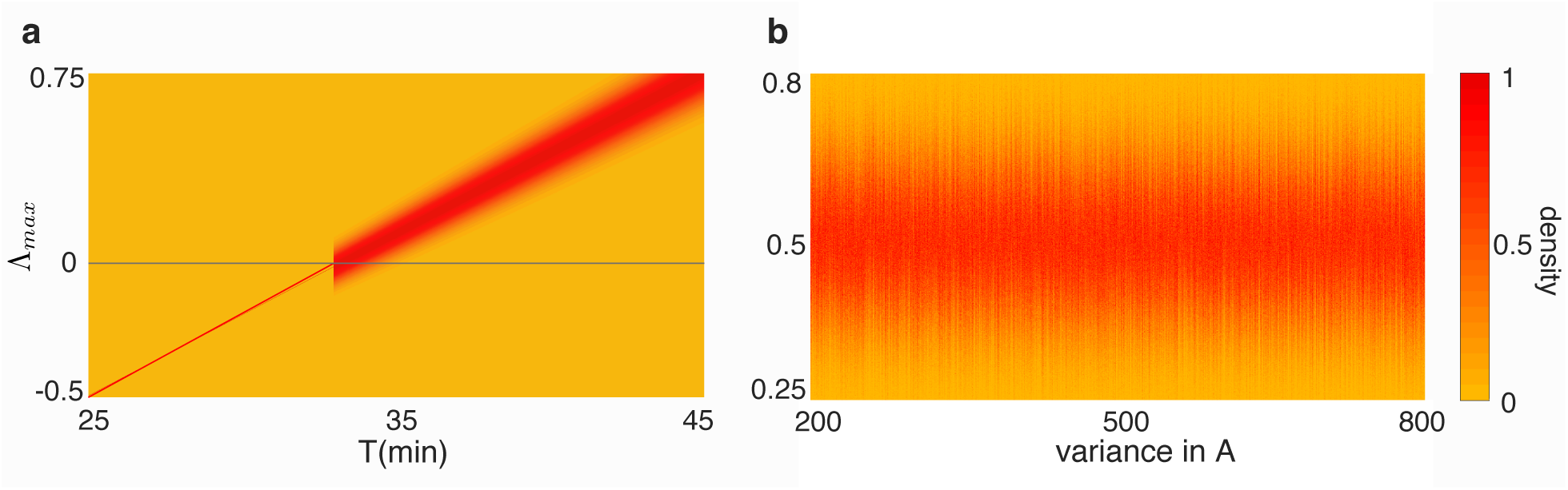
The distribution of Λ_max_ for the Ultradian model when the kicks are independent and uniformly distributed. Once again we see evidence of DIU. **(a)** We fix the kick amplitude distribution *A* ∼ *U* [45, 55], so that ⟨ *A*⟩ = 50 mg/dL and Var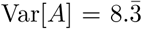. For *T* values that produce Λ_max_ *<* 0 in Figure 5c, the distribution of Λ_max_ resembles a Dirac delta. However, for *T* values that produce Λ_max_ > 0 in Figure 5c, a distribution of values emerges for Λ_max_ that broadens as *T* grows. **(b)** We fix *T* = 40 min, fix ⟨*A*⟩ = 50 mg/dL, and plot the distribution of Λ_max_ as a function of Var[*A*].

For our final experiment, we fix *T* at *T* = 40 min, fix the mean of the kick amplitude distribution at 50 mg/dL, and examine how the distribution of Λ_max_ varies with the variance of the kick amplitude distribution. Notice that *T* = 40 min is beyond the critical value at which we see an abrupt behavioral change in Figure 8a. Interestingly, the overall width of the Λ_max_ distribution seems to be insensitive to kick amplitude variance, yet we see subtle variation at fine scales (Figure 8b).

## 6 Discussion

### 6.1. Summary

In this paper we have proposed and analyzed a novel route by which delay can cause reliability failure in externally forced dynamical systems. The DIU recipe consists of three ingredients. First, delay must induce excitability in the unforced system. Second, the unforced system must possess shear. Finally, the external forcing must interact with the shear. We have shown that this recipe can transform limit cycles into strange attractors.

Our detailed analysis of delay linear shear flow illuminates the geometry of DIU and explicitly captures the role that delay plays in the onset of reliability failure. Building on this analysis, we have demonstrated that DIU exists in the glucose-insulin Ultradian model. Because many physiological systems oscillate and possess meaningful sources of delay, we believe DIU is relevant throughout mathematical physiology.

### 6.2. DIU in clinical medicine: Impact and outlook

Our results are directly applicable to ICU glucose care. The existence of delay-induced uncertainty in the Ultradian model and likely other models with delay acknowledges and potentially explains the difficulties clinicians have managing glucose in the ICU. It may feel to clinicians that if they just tried harder or had a better protocol or even had a patient-specific protocol, they would choose better treatments. If the process is chaotic, however, then predicting the precise evolution of the system becomes next to impossible. Recently, artificial intelligence has been touted as a potential solution to problems in health care [62]. Our results imply that artificial intelligence may not be the solution because glucose-insulin metabolism is not just complicated but actually chaotic. A better path might be to acknowledge the chaos and think statistically, estimating expected variances so that decisions can be based on the distribution of likely outcomes. Sampling more frequently can better track deviations and allow for interventions before dangerous glucose levels are reached. Better matching of models to real ICU experience may help identify and avoid interventions that are most likely to produce chaotic behavior. For example, smaller, more frequent insulin doses may be better than larger, less frequent doses. In the language of this paper, the former corresponds to lower kick amplitude and smaller inter-kick time.

Beyond the ICU, we believe DIU broadly impacts clinical decision-making. When treating a patient, a clinician prescribes a medical intervention, or lack thereof, based on experience: the outcome of randomized clinical trials, observational studies, or heuristic understanding. In many cases clinicians are not afforded the luxury of quantitative bases for decisions. Patient treatment is based on results of trial-and-error: An intervention is prescribed, and, if it fails, a new intervention is prescribed based on the results. However, such clinical empiricism assumes reliability of the physiological process being treated. DIU is thus important for potentially elucidating inexplicable failures in treatments based on experience.

Practical medicine induces additional complexity we will analyze in future work, most notably *nonstationarity*. In this paper we have analyzed stationary models, meaning that model parameters do not change over time. In the Ultradian context, we have therefore held the overall health state of the patient fixed, since the Ultradian model parameters represent overall health state. This is reasonable for glycemic management in the ICU, given the relatively fast treatment timescales. Over longer timescales, however, we anticipate that nonstationarity will be relevant for DIU analysis. We have set up a dichotomy in this paper: Assuming stationarity, does response to an external stimulus meaningfully depend on the state of the system (glucose level, plasma insulin level, interstitial insulin level in the Ultradian case) at the time the stimulus arrives? If no, we have reliability. If yes, we will observe DIU. How does this dichotomy generalize to nonstationary systems?

The notion of reliability we employ in this paper concerns the precise temporal evolution of *individual trajectories*. However, dynamical systems may also be viewed through an observational lens: *What are the statistical properties of observables of the system?* We associate DIU with reliability failure, yet the underlying rank one dynamics possess strong statistical regularity with respect to observables. Whether DIU is a benefit or a hazard therefore may depend on selecting the appropriate point of view for a particular application. We present two explicit examples to illustrate this point: glycemic management of enterally fed patients in an intensive care unit and glycemic self-management for a person with type-2 diabetes (T2DM).

In the ICU, enteral feeding induces glycemic oscillations. Overall patient health state will generally be nonstationary (because interventions aim to improve overall health state), but we can assume stationarity over the timescales of interest. The physiology of the patient will not be normal, and glucose levels will likely be elevated. Clinical management will proceed as follows. First, clinicians will maximize caloric intake. The nutrition level will be adjusted to the maximal amount the patient will tolerate, which may vary by hour. Next, blood glucose will be controlled by administering insulin. The clinicians try to avoid deadly hypoglycemia while minimizing hyperglycemia. As such, they make decisions about how to manage blood glucose based on the *observed boundaries* of glycemic dynamics. Clinicians measure glucose about once every hour or two, unless it is very low, in which case they measure it more frequently. Returning to DIU, decisions are made based on stability of statistics (such as glucose means and variances). Intensivists want predictable and stable glucose boundaries; they do not care about the exact temporal evolution of individual orbits. For DIU to impact gycemic management in the ICU, its presence must alter the statistics in a way that changes insulin dosage. We think that this is possible: *Inducing DIU might make the variance of glucose dynamics smaller and more stable with respect to other perturbations*.

The T2DM scenario contrasts sharply with glycemic management in the ICU. Until late in the disease, one manages type-2 diabetes with diet, exercise, and non-insulin medication. This requires patients to measure blood glucose before and after activities such as eating a meal, in order to quantify activity impact and make subsequent decisions. Here, rational decision-making requires understanding the temporal evolution of individual glucose trajectories. Ideally, reliable return to equilibrium follows perturbations such as breakfast and subsequent snacks. In the presence of DIU, however, glucose readings potentially become uninterpretable. This would render self-management a nearly impossible task, and suggests that DIU may negatively impact T2DM patients.

Our results set the stage for important future work. We have shown in this paper that the DIU phenomenon has the potential to confound medical science. Said differently, we have identified the problem. Solving the problem begins with detection. We must link DIU to data-driven science. How do we detect DIU using experimental data? What mechanisms produce DIU, and do these mechanisms possess specific detectable statistical signatures?

DIU phenomenology can be subtle and complex. Indeed, we have argued that DIU can have negative or positive impact, even in the same general physiological setting. When and how should we deliberately induce DIU? How can we mitigate negative impacts?

### 6.3. Directions for future mathematical research

Our work raises several intriguing mathematical problems. First, can one develop a general theory of rank one maps that would apply to the flows on function spaces generated by delay differential equations? There exist two lines of attack. One could build a theory of rank one maps that applies directly to Banach spaces. Alternatively, one could combine existing rank one theory with invariant manifold techniques, as Lu, Wang, and Young have done for supercritical Hopf bifurcations in certain parabolic PDEs [40]. Second, the external forcing we have studied here has relatively simple temporal structure (periodic or Poissonian pulsing). What do more complex forcing signals produce in the DIU context? For instance, what if the pulses arrive in a temporally inhomogeneous way, say localized around breakfast, lunch, and dinner? What if the forcing assumes the form of a continuous-time stochastic process with a jump component, such as Lévy noise?

Nonstationary dynamical systems have received considerable attention over the past fifteen years. Here, the dynamical model itself varies in time. Glycemic management in the ICU fits naturally into this setting: Ultradian model parameters likely drift over time as overall patient health state slowly changes. Moreover, we do not know the statistics of this parameter variability. (If we had this statistical information, we would be in the setting of random dynamical systems.) Can one develop DIU theory for nonstationary dynamical systems?

It will be important to deepen the links between DIU and data-driven science. Can one develop methods to deduce the presence of DIU from experimental data? What impact does DIU have on data assimilation schemes? In particular, how does the fact that the Ultradian model exhibits DIU impact our ability to fit this model to ICU patient data?

Finally, do there exist additional routes to DIU? What happens when delay acts on multiple timescales?

## 7. Author contributions

BK performed the numerical simulations. All authors developed the analysis and wrote the paper.

## 8. Funding

This work has been partially supported by National Science Foundation grant DMS 1816315 (WO) and by National Institutes of Health grants R01 LM006910 (DA, GH), R01 LM012734 (DA, GH), and R01 GM117138 (BK, WO).

## Notes

### Competing Interest Statement

The authors have declared no competing interest.

